# Caspase-8 activity mediates TNFα production and restricts *Coxiella burnetii* replication during murine macrophage infection

**DOI:** 10.1101/2024.02.02.578698

**Authors:** Chelsea A. Osbron, Crystal Lawson, Nolan Hanna, Heather S. Koehler, Alan G. Goodman

## Abstract

*Coxiella burnetii* is an obligate intracellular bacteria which causes the global zoonotic disease Q Fever. Treatment options for infection are limited, and development of novel therapeutic strategies requires a greater understanding of how *C. burnetii* interacts with immune signaling. Cell death responses are known to be manipulated by *C. burnetii*, but the role of caspase-8, a central regulator of multiple cell death pathways, has not been investigated. In this research, we studied bacterial manipulation of caspase-8 signaling and the significance of caspase-8 to *C. burnetii* infection, examining bacterial replication, cell death induction, and cytokine signaling. We measured caspase, RIPK, and MLKL activation in *C. burnetii*-infected TNFα/CHX-treated THP-1 macrophage-like cells and TNFα/ZVAD-treated L929 cells to assess apoptosis and necroptosis signaling. Additionally, we measured *C. burnetii* replication, cell death, and TNFα induction over 12 days in RIPK1-kinase-dead, RIPK3-kinase-dead, or RIPK3-kinase-dead-caspase-8^-/-^ BMDMs to understand the significance of caspase-8 and RIPK1/3 during infection. We found that caspase-8 is inhibited by *C. burnetii*, coinciding with inhibition of apoptosis and increased susceptibility to necroptosis. Furthermore, *C. burnetii* replication was increased in BMDMs lacking caspase-8, but not in those lacking RIPK1/3 kinase activity, corresponding with decreased TNFα production and reduced cell death. As TNFα is associated with the control of *C. burnetii*, this lack of a TNFα response may allow for the unchecked bacterial growth we saw in caspase-8^-/-^ BMDMs. This research identifies and explores caspase-8 as a key regulator of *C. burnetii* infection, opening novel therapeutic doors.

## Introduction

*Coxiella burnetii* (*C. burnetii*) is a Gram-negative, obligate intracellular bacterial pathogen and the causative agent of the global zoonotic disease query (Q) fever, also known as coxiellosis (1–3). Patients who contract Q fever typically present acute symptoms including fever, fatigue, and muscle aches; however, chronic disease can result in endocarditis and chronic fatigue syndrome, especially in those who are immunocompromised or pregnant (4). There is no widely available vaccine for *C. burnetii*, and current treatments involve antibiotic regimens which can last for over 18 months in chronic cases (5). Additionally, *C. burnetii* exhibits great environmental stability due to its small-cell variant’s (SCV) resistance to temperature and desiccation (3). This, combined with its ability to become aerosolized, has led to the labeling of *C. burnetii* as a potential bioterrorism threat (6, 7) and for the United States Center for Disease Control and Prevention (CDC) to categorize *C. burnetii* and select agent (8).

*C. burnetii* can infect a wide variety of animal hosts, including important livestock species such as sheep, cattle, and goats, as well as ticks, birds, and reptiles. Infected livestock animals are often asymptomatic, but suffer from spontaneous abortions, stillbirths, and weak offspring due to infection (9). Livestock also act as a reservoir from which bacteria is spread to humans, primarily through inhalation of bacteria from contaminated animal urine, feces, blood, milk, and birth products – the last of which has been shown to have high concentrations of bacteria (3, 10, 11). Outbreaks of Q fever in livestock populations have had significant economic consequences. Most well-known is the 2007-2011 Netherlands epidemic, during which disease on dairy goat farms led to estimated costs of 250-600 million euros associated with the over 4,000 human cases and the necessity for large-scale control measures including culling of pregnant animals, breeding restrictions, and strict monitoring of dairy products (12–15).

In general, *C. burnetii* is known to be largely immunologically silent, avoiding or suppressing innate immune signaling during infection (16). Bacterial effectors are vital for sustained suppression of immune signaling and survival of *C. burnetii* within the cell (17–21). Nevertheless, a recent report by Case *et al.* demonstrated that infection of primary murine bone-marrow-derived macrophages with *C. burnetii* elicits incomplete macrophage M1 polarization and decreased cytokine production during initial stages of infection independently of effector protein secretion (22), highlighting the diverse strategies that *C. burnetii* employs to manipulate its host cell environment.

This lack of a robust immune response can partially be attributed to the bacteria’s lifecycle within the cell, as it rapidly forms and replicates within a lysosomal-like compartment called the *Coxiella*-containing vacuole (CCV) (23) shielding its LPS from host immune sensors. While *C. burnetii* can infect a wide range of phagocytic and non-phagocytic cell types including HeLa cells, L929 cells, and macrophages, it has been demonstrated to preferentially infects alveolar macrophages in the lung environment (24–26). Infection of macrophages begins with internalization of the more metabolically silent and resilient SCV into phagosomes by α_V_β_3_ integrins (27), followed by the establishment of the CCV through fusion with endosomes, lysosomes, and phagosomes (23). As the pH within the CCV becomes more acidic, *C. burnetii* transitions from its SCV to its more metabolically active and replicative large-cell variant (LCV) (28–32). At this point, the bacteria’s Dot/Icm type IVB secretion system (T4SS) begins producing and secreting effector proteins from the CCV, of which over 140 candidates have been identified (33, 34). These effectors manipulate an array of host signaling pathways including, but not limited to, apoptosis, autophagy, inflammasome activation, transcription, and translocation (35). After approximately six days of growth within the cell, *C. burnetii* begins transitioning back into its SCV, such that both SCV and LCV can be found within the large CCV (29). At this point, the bacteria can spread to other cells through egress methods that have until recently been unknown, but are now known to involve host cell apoptosis (36).

Programmed cell death signaling, once limited to the ideas of apoptosis and necrosis (37), is now a complex area of research encompassing a plethora of other modes of regulated cell death such as necroptosis, pyroptosis, ferroptosis, and PANoptosis, the latter of which highlighting that the path to cell death is not a straight line and that signaling molecules in these pathways often have multiple roles (38, 39). For the purposes of this work, however, we will focus on apoptosis and necroptosis. Apoptosis is non-lytic, non-inflammatory, and can be divided into two types: intrinsic and extrinsic (40). Canonically, the intrinsic, or mitochondrial, pathway is triggered by cellular stress such as ROS, ER stress, or UV damage, and is regulated by mitochondrial release of cytochrome c and activation of caspase-9 (41–43). Induction of the extrinsic pathway, in contrast, is the result of activation of death receptors such as tumor necrosis factor alpha (TNFα) receptors (TNFRs) (40, 44). This leads to recruitment of death-inducing signaling complex (DISC) components to the TNFRs, notably TNFR-associated death domain (TRADD), TNFR-associated factors (TRAFs), receptor-interacting protein kinase 1 (RIPK1), and caspase-8 (45–48). This in turn leads to caspase-8 activation. Following caspase-9 or -8 activation, the two pathways converge, as both caspases activate caspase-3 to bring about apoptosis (49–51).

Not only does caspase-8 induce apoptosis, but it also inhibits necroptosis (52, 53). Apoptosis and necroptosis are, in some ways, opposing modes of programmed cell death (54). Apoptosis, on the one hand, is a slower, non-lytic, and non-inflammatory process; on the other hand, necroptosis is rapid, lytic, and highly inflammatory. However, both forms of cell death can be activated by TNFα. Specifically, if, following TNFR activation, caspase-8 is inhibited such that extrinsic apoptosis cannot occur, RIPK1 can interact with RIPK3 leading to phosphorylation of pseudokinase mixed lineage kinase domain-like protein (MLKL) (55–57). Phosphorylated MLKL (p-MLKL), then causes pore formation in the cell membrane and subsequent cell lysis (58).

Past work regarding host cell death during *C. burnetii* infection has primarily focused on intrinsic apoptotic signaling mediated by caspase-9 and the mitochondria and what methods the bacteria uses to subvert and control it (59). Indeed, several effector proteins have been identified which inhibit intrinsic apoptosis during early infection (60–66), and bacterial activation of apoptosis at late stages of infection has been documented (36). In contrast, while it has been shown that *C. burnetii* prevents extrinsic, caspase-8-mediated apoptosis induction at early stages of infection (67), the mechanisms by which this inhibition is accomplished and the significance to infection remain unknown. To address the knowledge gap surrounding bacterial interactions with cell death beyond intrinsic apoptosis, we investigated the interactions between *C. burnetii* and caspase-8. The focus of this study is on bacterial manipulation of caspase-8 signaling and the role of caspase-8 during *C. burnetii* infection, with an emphasis on consequences for bacterial replication, cell death induction, and cytokine signaling.

## Results

### *C. burnetii* inhibits TNFα-mediated caspase-8 activation

While *C. burnetii* has been documented to inhibit both intrinsic and extrinsic apoptosis, mechanistic details have only been investigated for bacterial manipulation of the intrinsic pathway (68). To determine if *C. burnetii* inhibits extrinsic apoptosis at the caspase-8 level, we treated *C. burnetii*-infected THP-1 macrophage-like cells with TNFα and cycloheximide (CHX) to induce extrinsic apoptosis. Briefly, differentiated THP-1 cells (seeded at 10^6^ cells/well in 12 well plates) were incubated with mCherry-expressing NMII *C. burnetii* (mCherry-*C. burnetii*) at a multiplicity of infection (MOI) of 25 GE/cell for 24 h, then washed to remove non-internalized bacteria (1 day post infection). At 3 days post infection (DPI), cells were pre-treated with 10 μg/mL CHX for 4 h, then incubated for 16 h with 20 ng/mL human TNFα.

Morphologically, *C. burnetii*-infected cells had an enlarged appearance consistent with harboring large CCVs and showed abundant mCherry signal (Fig 1A). While both infected and uninfected cells treated with TNFα/CHX visually had some cells that appeared to be dead or dying, by western blot it was obvious that *C. burnetii* infection had an inhibitory effect on cell death. Namely, we found that *C. burnetii* significantly reduce the cleavage of caspase-9, caspase-8, caspase-3, and PARP following TNFα treatment (Fig 1B). Previous work has demonstrated bacterial-inhibition of caspase-9, caspase-3, and PARP; however, this is the first report to our knowledge of caspase-8 inhibition by *C. burnetii* during extrinsic apoptosis or otherwise.

**Figure 1.**
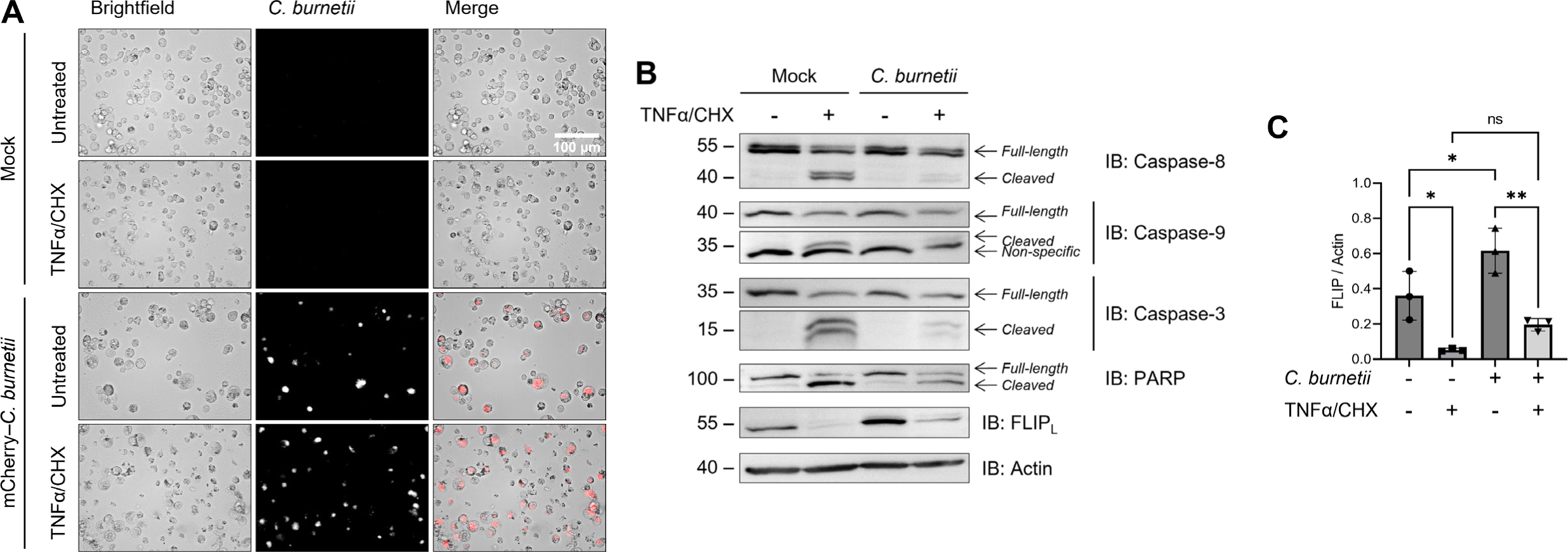
*C. burnetii* inhibition of caspase-8 activation and extrinsic apoptosis. THP-1 cells were differentiated using 100 nM PMA and infected with an mCherry-expressing Nine Mile Phase II (NMII) strain of *C. burnetii* (mCherry-*C. burnetii*) at MOI 25 GE/cell. At 3 DPI, cells were pre-treated with 10 μg/mL CHX for 4 h followed by overnight treatment with 20 ng/mL TNFα. (A) Representative images taken at 40x magnification. (B) Western blotting of samples as indicated. (C) Densitometry was completed in ImageJ and significance was determined by one-way ANOVA with Šίdák’s multiple comparisons test. Data are representative of three biological replicates from three independent experiments. *p < 0.05, **p < 0.01, ***p < 0.001, ****p < 0.0001

Furthermore, there was a marked upregulation of FLIP_L_ protein levels in our infected cells (Fig 1B-C), correlating with findings by Voth *et al.* regarding *cflip* mRNA upregulation (67). As FLIP is an inhibitor of caspase-8, this upregulation could be one way that *C. burnetii* inhibits caspase-8. Nevertheless, there was a significant reduction in FLIP levels following TNFα treatment regardless of infection, rendering it possible, if not likely, that FLIP upregulation is not the only method *C. burnetii* utilizes to prevent caspase-8 activation.

### *C. burnetii* infection sensitizes L929 cells to necroptosis

Caspase-8 is not only an initiator of apoptosis, but also a vital inhibitor of necroptosis. Indeed, the knockout of caspase-8 in mice is embryonically lethal due to an over-activation of RIPK3 leading to necroptosis (69). Therefore, by inhibiting caspase-8, *C. burnetii* could inadvertently trigger necroptotic signaling during infection. To investigate this possibility, we infected L929 cells (seed at 5×10^4^ cells/well in 24 well plates) with mCherry-*C. burnetii* at an MOI of 600 GE/cell, incubating cells with bacteria for 24 h then washing to remove non-internalized bacteria (1 DPI). As L929 cells are non-phagocytic, we observed a much lower rate of infectivity than in our macrophage cells, which is what led to the usage of a higher MOI here. At 3 and 6 DPI, L929 cells were treated with the pan-caspase inhibitor Z-VAD-FMK (ZVAD) for 30 minutes, followed by treatment with mouse TNFα for 3 h.

Phenotypically, both infected and mock-infected cells at 3 and 6 DPI which were treated with TNFα/ZVAD appeared rounded and swollen, consistent with necroptosis (Fig 2A and D). Additionally, we found that the phosphorylation of RIPK1, RIPK3, and MLKL in response to TNFα/ZVAD treatment was amplified in our infected cells at both 3 (Fig 2B) and 6 DPI (Fig 2E). Indeed, in the case of MLKL, densitometry analysis revealed that this phosphorylation was approximately 1.5-fold and 3-fold higher in *C. burnetii*-infected cells at 3 and 6 DPI, respectively (Fig 2C and F). These data indicate that bacterial infection intensifies necroptosis in L929 cells, notably more at 6 DPI than at 3 DPI, suggesting a relationship between the stage of infection and the sensitivity of infected cells to necroptosis.

**Figure 2.**
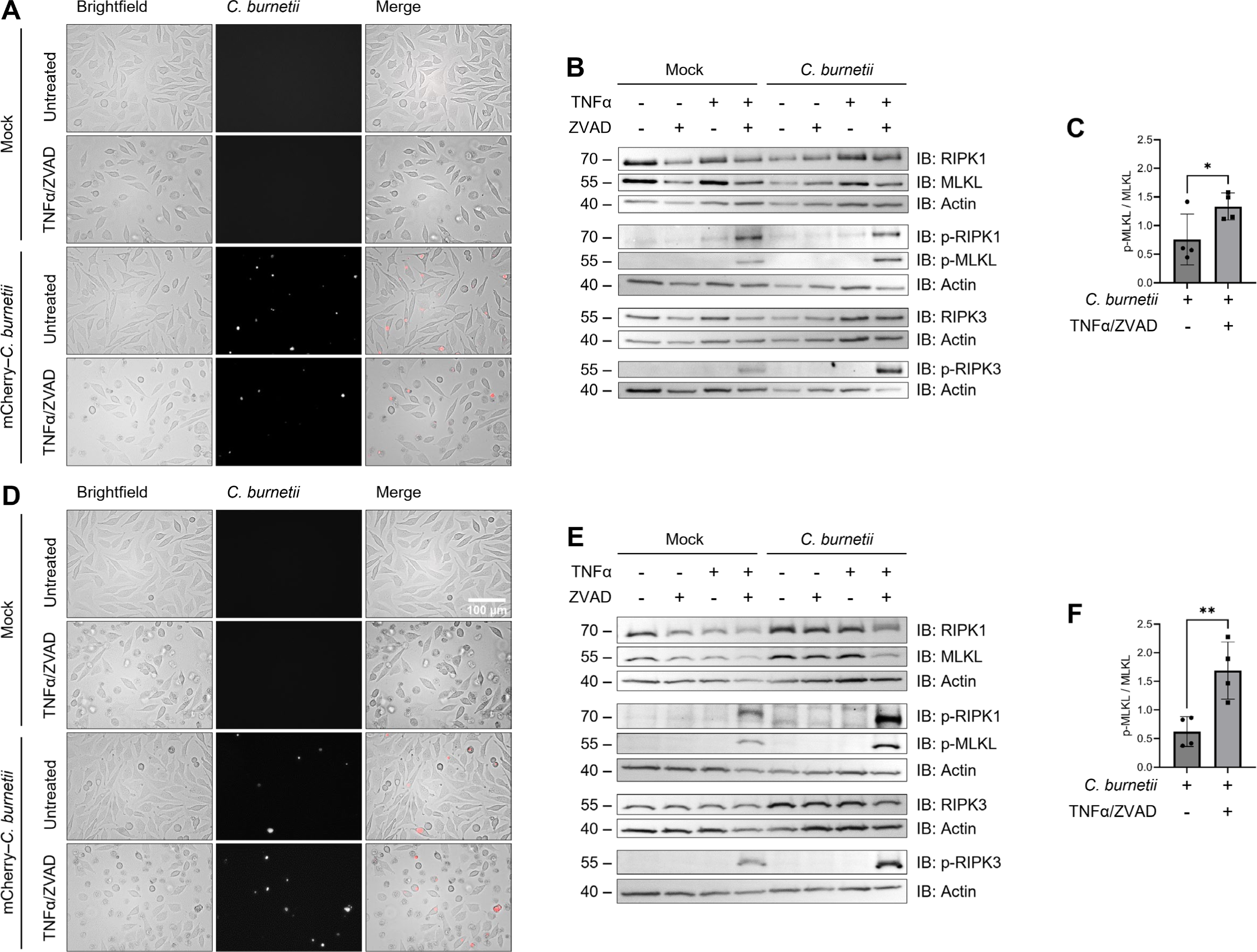
Necroptosis exacerbation by *C. burnetii* infection. L929 cells were infected with mCherry-*C. burnetii* at MOI 300 GE/cell. At 3 and 6 DPI, cells were pre-treated with 50 μM Z-VAD-FMK for 30 mins followed by 3 h incubation with 20 ng/mL TNFα. (A,D) Representative 3 and 6 DPI images taken at 40x magnification. (B,E) Western blotting of samples as indicated. (C,F) Densitometry was completed in ImageJ and significance was determined by paired t-test. Data are representative of four biological replicates from four independent experiments. *p < 0.05, **p < 0.01, ***p < 0.001, ****p < 0.0001

Interestingly, we neither observed dying cells nor detected phosphorylation of RIPK1, RIPK3, or MLKL in our infected cells treated with TNFα alone (Fig 2B), indicating that the caspase inhibition by *C. burnetii* is not as potent as our pharmacological inhibition and may not be substantial enough to result in cell death during typical cellular conditions. Thus, we concluded that *C. burnetii* infection sensitizes L929 cells to necroptosis when induced but does not appear to induce it on its own at these timepoints.

### *Ripk3^K51A/K51A^Casp8^-/-^* BMDMs have increased *C. burnetii* replication at late stages of infection

Thus far, we have documented for the first time that *C. burnetii* (1) inhibits caspase-8 to prevent extrinsic apoptosis in THP-1 cells and (2) sensitizes L929 cells to necroptosis. Nevertheless, it remains to be seen if caspase-8 or RIPK activity is an important aspect of the immune response to *C. burnetii* infection outside of situations of pharmacological induction. To determine whether caspase-8, RIPK1, or RIPK3 activity restrict *C. burnetii* replication, we utilized BMDMs derived from femurs of C57BL/6J wild-type (WT), *Ripk1^K45A/K45A^* (RIPK1-kinase-dead, R1KD) (70), *Ripk3^K51A/K51A^* (RIPK3-kinase-dead, R3KD) (71), and *Ripk3^K51A/K51A^Casp8^−/−^*(RIPK3-kinase-dead-caspase-8^-/-^, R3KDCasp8^-/-^) (71) mice. Notably, these kinase-dead BMDMs are still able to produce RIP1 and RIP3 proteins, but those kinases are unable to function enzymatically. Furthermore, since the knockout of caspase-8 on its own is embryonically lethal in mice (69), both WT and R3KD BMDMs serve as controls for the R3KDCasp8^-/-^ BMDMs.

We infected these BMDMs (seeded at 10^5^ cells/well in 12 well plates) with mCherry-*C. burnetii* at an MOI of 300 GE/cell by incubating cells with bacteria for 1 hour in media containing 2% FBS, followed by thorough washing to remove non-internalized bacteria (0 DPI). To facilitate infection, cells and bacteria were centrifuged at 300 rcf for 10 minutes at the start of the 1 hour incubation. To quantify bacteria load, we measured bacterial genome equivalents (GE) at 3, 6, 9, and 12 DPI via quantitative real-time polymerase chain reaction (qPCR) targeting the *C. burnetii dotA* gene.(29) To further probe the course of infection in these cells, we also assessed the percentage of cells which were infected. For these experiments, BMDMs were seeded at 2 x 10^4^ cells/well in 96 well plates and were infected as described above with mCherry-*C. burnetii*. At 6 and 12 DPI, cells were stained with Hoechst to label nuclei and the percentage of cells which were also mCherry-positive was quantified.

*C. burnetii* was able to productively infect and replicate within all the genotypes tested (Fig 3A), and our qPCR analysis revealed that, while bacterial load was not different during early stages of infection, by 12 DPI there was significantly more bacteria in BMDMs lacking caspase-8 than in any other genotype (Fig 3B). In fact, we detected a ∼2.5-fold increase in the amount of *C. burnetii* in the infected R3KDCasp8^-/-^ BMDMs compared to the WT and R3KD BMDMs, and a ∼1.5-fold increase in the amount compared to the R1KD BMDMs. This increased replication in R3KDCasp8^-/-^ BMDMs was accompanied by an approximately 2-fold increase in the percent of cells which harbored bacteria compared to all other genotypes (Fig 3C). These results support the role of caspase-8 in restricting *C. burnetii* replication and spread, specifically during late-stage infection. Moreover, as there was not higher bacterial load in the R1KD and R3KD cells compared to the WT, it is unlikely that necroptosis is a key regulator of NMII *C. burnetii* replication in BMDMs.

**Figure 3.**
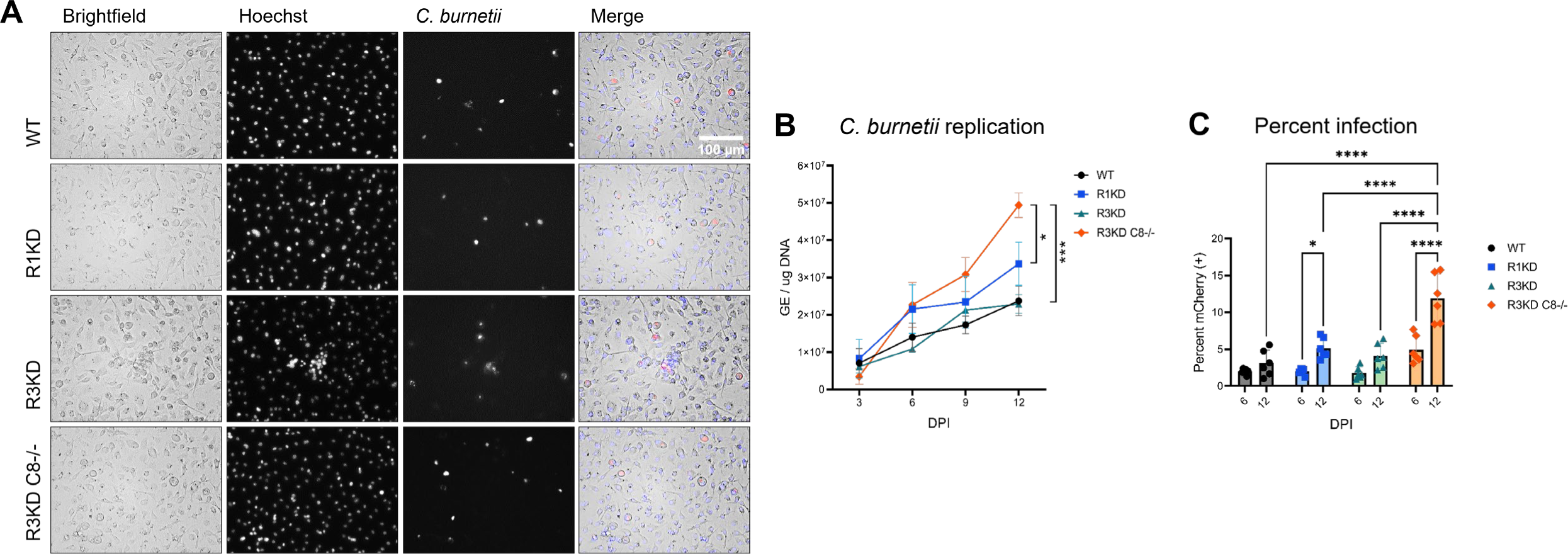
Caspase-8 restricts *C. burnetii* replication and spread. Primary murine bone marrow-derived macrophages (BMDMs) (C57B6J WT, *Ripk1^K45A/K45A^*, *Ripk3^K51A/K51A^*, and *Casp8^-/-^ Ripk3^K51A/K51A^*) were infected with mCherry-*C. burnetii* at MOI 100 GE/cell. At 3, 6, 9, and 12 DPI, cells were imaged and lysed using a MP Biosciences FastPrep-24 machine and 0.1 mm zirconia beads. (A) Representative 12 DPI images taken at 40x magnification. (B) Quantification of *C. burnetii* genome equivalents (GE) as determined by *DotA* qPCR performed on whole cell lysates. Significance was determined by mixed effects analysis with Tukey’s multiple comparisons test. Data are representative of three biological replicates from three independent experiments. Error bars in (B) represent SEM instead of SD. (C) Percent infection was measured at 6 and 12 DPI using a Molecular Devices ImageXpress Micro Confocal. Data represents 6 biological replicates, and 6 and 12 DPI datasets are from two independent experiments. Significance was determined by two-way ANOVA with Tukey’s multiple comparisons test. *p < 0.05, **p < 0.01, ***p < 0.001, ****p < 0.0001

### Cell death is reduced in *Ripk3^K51A/K51A^* and *Ripk3^K51A/K51A^Casp8^-/-^*BMDMs throughout *C. burnetii* infection

Caspase-8 has multiple roles within the cell and is a key regulator of cell death pathways including apoptosis, necroptosis, and pyroptosis (72). This, combined with the importance of cell death signaling pathways to *C. burnetii* infection (59), renders it possible that disruption of cell death regulation in the caspase-8-negative BMDMs is responsible for the increased susceptibility to infection. To determine if cell death levels were altered in R3KDCasp8^-/-^ BMDMs during *C. burnetii* infection, we assessed cytotoxicity by SYTOX staining. For these experiments, BMDMs were seeded and infected in 96-well plates as in Fig 3 with mCherry-*C. burnetii*. At 6 and 12 DPI, cells were stained with SYTOX to identify dying cells and stained with Hoechst to label nuclei, and the percentage of cells which were SYTOX-positive was calculated.

We found that, at 6 (Fig 4A) and 12 DPI (Fig 4B), both R3KD and R3KDCasp8^-/-^BMDMs had significantly reduced cytotoxicity compared to WT. In contrast, R1KD BMDMs only had reduced cytotoxicity compared to WT at 12 DPI. In the case of the R3KD cells, the decrease in cell death throughout infection is likely due to the lack of RIPK3-mediated necroptosis. While the R3KDCasp8^-/-^ BMDMs lack both RIPK3-mediated necroptosis and caspase-8-mediated apoptosis, the combined loss of both pathways unexpectedly did not amplify the loss of cytotoxicity. In the R1KD cells, while there is a lack of RIPK1-mediated necroptosis, necroptosis via other pathways should remain unaltered as RIPK3 is still active. Furthermore, though the scaffolding roles of RIPK1 in regulated apoptosis should be preserved in the R1KD cells, it is possible that some dysregulation of apoptosis occurs (73–75). Thus, the decreased cell death in the R1KD BMDMs at 12 DPI may indicate that RIPK1-mediated cell death is induced at late stages of infection. In contrast, we found that cell death due to infection was not dependent on RIPK1 kinase activity at 6 DPI. Overall, we concluded that the activity of RIPK1 and RIPK3 are both key factors in mounting a cell death response to *C. burnetii* infection in BMDMs, with RIPK1-mediated cell death playing a more important role at late stages of infection compared to early stages. Additionally, our results suggest that caspase-8 plays either a limited role in the cell death response to infection, or one that is dependent on RIPK3.

**Figure 4.**
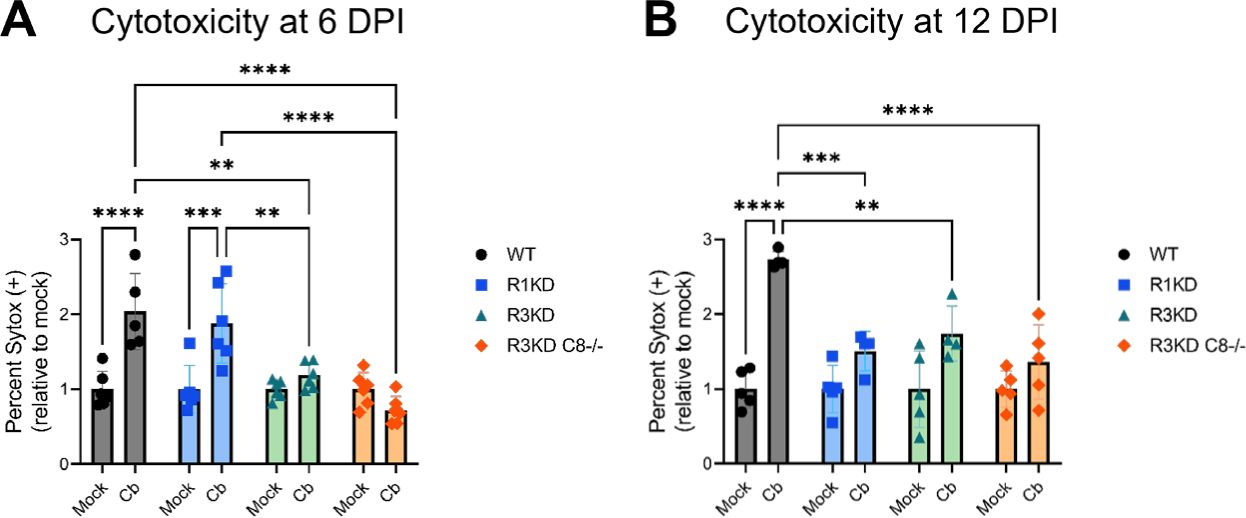
BMDMs lacking RIPK3 or RIPK3 and caspase-8 activity have decreased cytotoxicity throughout *C. burnetii* infection. BMDMs were infected with mCherry-*C. burnetii* as in Fig 3. At 6 DPI (A) and at 12 DPI (B), cytotoxicity was measured by SYTOX staining using a Molecular Devices ImageXpress Micro Confocal. Data are representative of four-six biological replicates from two independent experiments. Percent SYTOX positive was normalized to the average mock cytotoxicity within genotypes and significance was determined by two-way ANOVA with Tukey’s multiple comparisons test. *p < 0.05, **p < 0.01, ***p < 0.001, ****p < 0.0001

**Figure 5.**
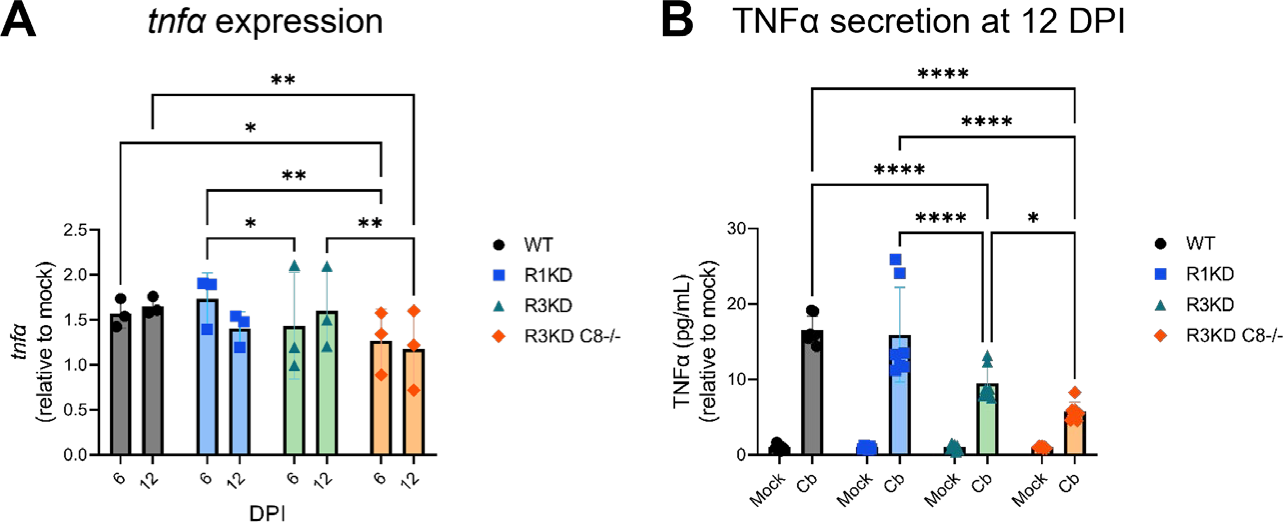
TNFα production is reduced in caspase-8^-/-^ BMDMs during *C. burnetii* infection. BMDMs were infected with mCherry-*C. burnetii* as in Fig 3. (A) Relative expression of *tnfa* at 12 DPI was determined by qRT-PCR. Ct values were normalized first to *gapdh* expression, then to mock-infected samples. Data are representative of three biological replicates from three independent experiments, and significance was determined by two-way ANOVA with Tukey’s multiple comparisons test. (B) Quantification of TNFα at 12 DPI in cell-free supernatant. TNFα pg/mL concentrations were normalized to the average mock values within genotypes. Data are representative of seven biological replicates from two independent experiments. Significance was determined by two-way ANOVA with Tukey’s multiple comparisons test. *p < 0.05, **p < 0.01, ***p < 0.001, ****p < 0.0001

### *Ripk3^K51A/K51A^Casp8^-/-^* BMDMs have an attenuated TNFα response to *C. burnetii* infection

Beyond apoptosis and necroptosis, caspase-8 plays a key scaffolding role during TNFα-mediated cytokine signaling (72). Therefore, to further discern what signaling changes are occurring in our R3KDCasp8^-/-^ BMDMs that could allow for increased *C. burnetii* growth, we investigated TNFα induction during infection by quantitative real-time reverse-transcription PCR (qRT-PCR) and enzyme-linked immunosorbent assay (ELISA). For qRT-PCR experiments, BMDMs were seeded at 10^5^ cells/well in 12 well plates and were lysed in TRIzol at 6 and 12 DPI. For ELISA experiments, BMDMs were seeded at 2 x 10^4^ cells/well in 96 well plates and cell-free supernatant was collected at 12 DPI. In both sets of experiments, BMDMs were infected as in Fig 3 with mCherry-*C. burnetii*.

Our qRT-PCR analysis showed a reduction of *tnfα* expression in infected R3KDCasp8^-/-^ BMDMs compared to WT at 6 DPI and an even greater reduction compared to both the WT and R3KD BMDMs at 12 DPI. Quantification of TNFα secretion via ELISA at 12 DPI supported this finding, where we found a moderate reduction in TNFα concentration in cell-free supernatant of R3KD BMDMs and a large reduction in R3KDCasp8^-/-^ cells. Together, these data indicate that, without caspase-8, BMDMs initiate a highly attenuated TNFα response to *C. burnetii* infection in BMDMs. As TNFα has been associated with restricting *C. burnetii* growth within cells (76–79), it is possible that this lack of TNFα production in R3KDCasp8^-/-^ BMDMs leads to increased bacterial replication in these cells.

## Discussion

In this study, we provide one of the first reports for the role of caspase-8 during *C. burnetii* infection. Using THP-1 and L929 infection models, we demonstrated that *C. burnetii* inhibits caspase-8 activation during TNFα-mediated apoptosis and sensitizes cells to TNFα-mediated necroptosis. We also showed that caspase-8 restricts *C. burnetii* replication, as BMDMs lacking caspase-8 showed increased bacterial load and percent infection by 12 DPI. The increased bacterial spread in caspase-8-negative cells is likely connected to the increased bacterial replication, as it was recently discovered that *C. burnetii* egress is dependent on bacterial load (36). This higher susceptibility of caspase-8 deficient BMDMs to *C. burnetii* during late stages of infection also corresponded with decreased TNFα production and lower cytotoxicity, though the latter appeared to also be tied to the loss of RIPK3 activity and is thus not likely to be the sole contributing factor. As TNFα has been associated with the control of *C. burnetii*, it is possible that the lack of a TNFα response in our caspase-8-negative cells allowed for the unchecked bacterial growth.

During infection, *C. burnetii* heavily manipulates the host cell environment to control apoptosis and several anti-apoptotic effector proteins have been identified (59). Until now, however, the ability of *C. burnetii* to inhibit caspase-8 was unknown, despite its key role in regulating not only apoptosis, but also necroptosis and pyroptosis (72). Previously, it was shown that TNFα-induced apoptosis is inhibited by *C. burnetii* (64, 67) and that the *C. burnetii* effector CaeA is able to prevent apoptosis at the executioner caspase-3 level (64), but whether bacterial manipulation of extrinsic apoptosis is limited to downstream steps or also includes upstream signaling had not been investigated. We have demonstrated that, indeed, caspase-8 activation and subsequent apoptosis is prevented by *C. burnetii* infection, extending past research and highlighting the extent to which *C. burnetii* is able to interfere with host cell signaling. Additionally, we were able to establish that the caspase-8 inhibitor cFLIP is over-expressed in *C. burnetii*-infected cells, validating and expanding findings by Voth *et al.* which showed increased cFLIP mRNA levels during infection (67). This over-expression likely contributes to bacterial inhibition of caspase-8, though it is also possible, if not probable, that effector proteins are involved, as with *C. burnetii* inhibition of other apoptotic caspases (60, 65, 64). An effector screen, as well as a mutant library screen, would be beneficial in determining which effector proteins or other bacterial factors, known or unknown, are involved in the inhibition of caspase-8.

As discussed earlier, caspase-8 sits at the nexus of multiple cell death pathways, and caspase-8 inhibition is a critical step in induction of necroptosis (54). Because of its inhibition of caspase-8, we theorized that cells infected with *C. burnetii* would be susceptible to necroptotic death. Indeed, we found that infected L929 cells treated to induce necroptosis had ∼3-fold more MLKL phosphorylation than uninfected cells at 6 DPI. This vulnerability to necroptosis is a vital piece of information, as it suggests a possible blind-spot in the anti-cell death regime of *C. burnetii*. It is possible that, similarly to its approach to pyroptosis and in contrast to its approach to apoptosis, *C. burnetii* primarily utilizes a stealth strategy as opposed to a defensive one to avoid necroptotic cell death (24, 78, 80–82). However, whether necroptosis results in the killing of bacteria within the cell or if it is a mechanism by which *C. burnetii* spread can occur remains to be seen. The effect of necroptosis on *C. burnetii* is likely also dependent on the stage of infection. If the CCV is predominately composed of LCV bacteria, for instance, necroptosis may have a higher bactericidal effect than if the CCV predominately contains the more resistant SCV bacteria. Surprisingly, though *C. burnetii* infection rendered L929 cells more vulnerable to necroptosis, loss of RIPK3 kinase activity, and thus the ability to undergo necroptotic cell death, did not result in a difference in susceptibility of BMDMs to *C. burnetii*. This is intriguing, as Ripk3^-/-^ BMDMs have recently been found to have higher bacterial loads by 7 DPI compared to WT BMDMs, a phenotype lost in Ripk3^-/-^Caspase-8^-/-^ BMDMs (83). These data, combined with our own bacterial replication data, suggest that RIPK3 has significance to combatting *C. burnetii* infection independent of its kinase activity but dependent on caspase-8.

Despite the apparent dispensability of RIPK1 and RIPK3 kinase activity to controlling *C. burnetii* replication, we did see an approximately 2-fold and 3-fold increase in cell death by 6 and 12 DPI, respectively, in infected WT BMDMs which was dependent on RIPK1 activity at 12 DPI and on RIPK3 activity at both 6 and 12 DPI. These results implicate necroptotic machinery in cell death from infection. Overall, our bacterial replication and cell death assays suggest that, while necroptotic signaling components are involved in cell death during infection, they are dispensable to controlling or supporting *C. burnetii* replication.

Interestingly, our data also suggest that RIPK1 may have more involvement in cell death at late stages of infection, possibly in relation to increased (but still moderate) TNFα secretion at these time points. Additionally, while RIPK1 kinase activity is well-known as an inducer of necroptosis alongside RIPK3, the kinase activity of RIPK1 has also been associated with apoptosis regulation (73–75). Thus, it is possible that multiple cell death pathways are dysregulated in the R1KD cells. This nuanced role of RIPK1 in cell death at different timepoints during *C. burnetii* infection should be further probed to assess if and how its activity could be leveraged therapeutically.

The combined loss of caspase-8 and RIPK3 activity surprisingly did not result in a further decrease to cell death, despite our observed importance of caspase-8 to TNFα production. The mechanisms by which TNFα is able to control *C. burnetii* infection is an ongoing area of research, with past work showing that TNFα is vital to IFNγ-mediated control of *C. burnetii* (76, 77), and more recent studies implicating TNFα in restricting bacterial replication following toll-like receptor (TLR) activation (78) and in hypoxic conditions (79). Mechanistically, Boyer *et al*. found that TNFα restriction of *C. burnetii* replication in BMDMs involves IRG1-itaconate signaling (83), a pathway which has also been implicated by Kohl *et al*. (84). While TNFα treatment appears to be able to control *C. burnetii* in cells lacking caspase-8,(83) this study adds caspase-8 to the mix of endogenous TNFα regulators during infection. Further characterizing the role that caspase-8 plays within the mechanisms of TNFα-mediated control of *C. burnetii*, for cell death signaling and otherwise, will enhance our ability to harness the innate immune system to fight infection.

Moreover, this work, combined with past research documenting the increased disease severity in *C. burnetii*-infected mice deficient in TNFα (85), raises the possibility that patients taking TNFα blockers may be particularly vulnerable to Q fever. Following the outbreak in the Netherlands, one group attempted to address this very question by examining *C. burnetii* seroprevalence and chronic disease in patients with rheumatoid arthritis (RA) who were on anti-TNFα therapy (86). While patients with RA had a much higher prevalence of chronic Q fever overall (87), convincing conclusions were unable to be drawn regarding the risk of anti-TNFα therapies in particular due to their limited sample size. Recently, the potential for selectively inhibiting or activating TNFR1 and TNFR2 to treat inflammatory and degenerative diseases as an alternative to broad inhibition of both receptors was reviewed by Fischer *et al*. (88). The differential roles of TNFR1 and TNFR2 in the context of *C. burnetii* infection have not, to our knowledge, been explored, though TNFR2 upregulation in monocytes has been associated with Q fever endocarditis (89). This line of research should be further pursued to conclusively determine the connection between anti-TNFα therapy and Q fever risk.

The importance of caspase-8 during pathogenic infection has been documented in the context of many pathogenic species, including *Yersinia* (90–93) and herpes simplex virus 1 (HSV1) (94–96). In the case of *Yersinia*, the effector protein YopJ inhibits transforming growth factor β-activated kinase 1 (TAK1) and IKKβ (97–100), leading to RIPK1-dependent induction of caspase-8-mediated cleavage of gasdermin D (GSDMD) to induce pyroptosis and secretion of IL-1β (90–93). Our investigation of the relationships between caspase-8 and *C. burnetii* is a vital step in the path to understanding the significance of programmed cell death signaling to not only *C. burnetii* infection, but also to other pathogens, obligate intracellular bacteria or not, which interact with these pathways. As such, a deeper exploration of the mechanisms behind our findings will aid in the development of novel therapeutic strategies to improve animal and human health.

## Methods Cell Culture

Human THP-1 monocytes were maintained at a density between 3×10^5^ and 10^6^ cells/mL in 1x RPMI 1640 (Gibco 11875093) supplemented with 10% FBS (Cytiva Hyclone SH30070.03HI), 1 mM sodium pyruvate (Gibco 11360070), 10 mM HEPES (Gibco 15630106), and 50 μM beta-mercaptoethanol (Gibco 21985023), and 1x antibiotic-antimycotic (Gibco 15240062) at 37°C in 5% CO_2_. For differentiation into macrophage-like cells using phorbol 12-myristate 13-acetate (PMA; Sigma-Aldrich P1585), THP-1 monocytes were seeded in media containing 100 nM PMA. After 24 h, PMA-containing media was replaced with media containing no PMA, and cells were allowed to rest for 24 h prior to infection.

Mouse L929 cells were maintained in 1x DMEM, high glucose (Gibco 11965092) supplemented with 10% FBS (Atlas Biologicals EF-0500-A) and 1x antibiotic-antimycotic (Gibco 15240062) at 37°C in 5% CO_2_.

Primary murine bone marrow macrophages (BMDMs) were derived from bone marrow cells isolated from femurs of C57BL/6J WT, *Ripk1^K45A/K45A^*, *Ripk3^K51A/K51A^*, and *Ripk3^K51A/K51A^Casp8^−/−^*mice (70, 71). Briefly, bone marrow cells were differentiated for 7-10 days using 1x DMEM, low glucose, pyruvate (Gibco 11885084) supplemented with 10% FBS (Cytiva Hyclone SH30070.03HI), 30% L929 conditioned media (LCM), and 1x antibiotic-antimycotic (Gibco 15240062) at 37°C in 5% CO_2_, and 1/2 volume of media was replaced every 3 days.

### Bacterial stock and infection

mCherry expressing *C. burnetii* NMII (clone 4 RSA439) (mCherry-*C. burnetii*) was grown in Acidified Citrate Cysteine Medium 2 containing tryptophan (ACCM-2 + tryptophan) as previously described (101–103). *C. burnetii* bacterial stocks were quantified by quantitative polymerase chain reaction (qPCR) to measure genome equivalents (GE) (29, 104).

Differentiated THP-1 cells were washed 2x with incomplete 1x RPMI 1640 (Gibco 11875093) then incubated with mCherry-*C. burnetii* at a MOI of 25 GE/cell for 24 h in 1x RPMI 1640 supplemented with 10% heat-inactivated FBS (Cytiva Hyclone SH30070.03HI), 1 mM sodium pyruvate (Gibco 11360070), 10 mM HEPES (Gibco 15630106), and 50 μM beta-mercaptoethanol (Gibco 21985023). Following infection, THP-1 cells were washed twice with incomplete 1x RPMI 1640 to remove extracellular bacteria (1 DPI) and maintained in 1x RPMI 1640 supplemented with 10% FBS, 1 mM sodium pyruvate, 10 mM HEPES, and 50 μM beta-mercaptoethanol, with 1/2 volume of media replaced daily.

L929 cells were washed 2x with incomplete 1x DMEM, high glucose (Gibco 11965092) then incubated with mCherry-*C. burnetii* at a MOI of 600 for 24 h in 1x supplemented with 10% FBS (Atlas Biologicals EF-0500-A) with a 30 minutes spin at 300 rcf. Following infection, L929 cells were washed 2x with incomplete 1x DMEM to remove extracellular bacteria (1 DPI) and maintained in 1x DMEM supplemented with 10% FBS.

Differentiated BMDMs were washed twice with incomplete 1x DMEM, low glucose, pyruvate (Gibco 11885084) then incubated with mCherry-*C. burnetii* at a MOI of 300 GE/cell for 1 h in 1x DMEM supplemented with 2% FBS (Cytiva Hyclone SH30070.03HI) with a 10 minutes spin at 300 rcf. Following infection, BMDMs were washed twice with incomplete 1x DMEM to remove extracellular bacteria (0 DPI) and maintained in 1x DMEM supplemented with 10% FBS and 10% LCM, with 1/2 volume of media replaced daily.

### Cell Death Induction

For extrinsic apoptosis induction in THP-1 macrophage-like cells, cells were pre-treated with 10 μg/mL cycloheximide (CHX; Selleckchem S7418) for 4 h, then treated with 20 ng/mL human tumor necrosis factor alpha (TNFα; Sigma-Aldrich H8916) for 16 h.

For necroptosis induction in L929 cells, cells were pre-treated with 50 μM Z-VAD-FMK (Calbiochem/Sigma-Aldrich 627610) for 30 minutes followed by 3 h treatment with 20 ng/mL mouse TNFα (Calbiochem/Sigma-Aldrich 654245).

### Immunoblotting

Protein extracts were prepared either by lysing cells directly in 2x Laemmli buffer (0.125 M Tris-HCL pH 6.8, 4% SDS w/v, 20% glycerol, 0.004% bromophenol blue w/v, 10% beta-mercaptoethanol) (for necroptosis experiments in L929 cells and BMDM experiments) or by lysing cells in RIPA buffer (25 mM Tris-HCl pH 7.6, 150 mM NaCl, 1 mM EDTA, 1% NP-40, 1% sodium deoxycholate, 0.1% SDS, 1mM Na3VO4, 1 mM NaF, 0.1 mM PMSF, 10 μM aprotinin, 5 μg/mL leupeptin, 1 μg/mL pepstatin A) followed by dilution in 2x Laemmli buffer (for extrinsic apoptosis experiments in THP-1 cells). After mixing with Laemmli buffer, samples were heated for 5-10 minutes at 95°C and centrifuged for 5 minutes at 12,000 rcf. Samples were separated by SDS-PAGE using 10% or 15% acrylamide gels, followed by transfer onto 0.45 μm PVDF membranes (Immobilon/Sigma-Aldrich IPVH00010). Membranes were blocked for 30 minutes at room temperature with 5% nonfat dry milk (Lab Scientific/VWR M0841) or 5% BSA (ThermoFisher BP9706) in Tris-buffered saline (50 mM Tris-HCl pH 7.5, 150 mM NaCl) containing 0.1% Tween-20 (ThermoFisher BP337). Membranes were then labeled with primary antibody (see detailed list of antibodies below) overnight rocking at 4°C, then labeled with either anti-rabbit IgG-HRP conjugate (1:10,000; Promega W401B) or anti-mouse IgG-HRP conjugate (1:10,000; Promega W402B) for 2 h rocking at room temperature. Blots were visualized using an Analytik Jena UVP ChemStudio and either SuperSignal West Pico PLUS Chemiluminescent Substrate, SuperSignal West Femto Chemiluminescent Substrate, or SuperSignal West Atto Chemiluminescent Substrate (ThermoFisher PI34580, PI34096, PIA38554).

Primary antibodies used, all at 1:1000 unless otherwise specified: anti-caspase-3 (Cell Signaling 9662), anti-cleaved-caspase-3 (Cell Signaling 9661), anti-caspase-8 (Cell Signaling 9746), anti-caspase-9 (Cell Signaling 9504), anti-PARP (Cell Signaling 9542), anti-FLIP (Cell Signaling 56343), anti-RIPK1 (Cell Signaling 3493), anti-phosphorylated-RIPK1 (Cell Signaling 53286), anti-RIPK3 (Cell Signaling 15828), anti-phosphorylated-RIPK3 (Cell Signaling 91702), anti-MLKL (Cell Signaling 37705), anti-phosphorylated-MLKL (Abcam ab196436), anti-actin (1:10,000; Sigma-Aldrich A5441).

### Bacterial genome equivalents (GE) assay

To determine the bacterial load in BMDMs, cells were collected in 300 μL 1x PBS per 5 x 10^5^ cells and transferred to 1.5 mL screw-cap microtubes containing 100 μL of 0.1 mm zirconia/silica beads (BioSpec Products 11079101z). Bacterial DNA was released by homogenization three times at 5.0 m/s for 30 seconds using an MP Biosciences FastPrep-24, followed by centrifugation for 1 minute at 12,000 rcf. DNA was quantified by spectrophotometer (Biotek Cytation 3), and 50 ng of DNA diluted in molecular grade water was used for qPCR quantification targeting the *C. burnetii dotA* gene.(29)

### Percent infection assay

BMDMs were seeded at a density of 1×10^4^ cells/well in 96-well plates and infected with mCherry-*C. burnetii* at an MOI of 100 GE/cell as described above. At 6 and 12 DPI, cells were stained with Hoechst 33342 (2’-[4-ethoxyphenyl]-5-[4-methyl-1-piperazinyl]-2,5’-bi-1H-benzimidazole) (Biorad 1351304). Using a Molecular Devices ImageXpress Micro Confocal, total number of cells (Hoechst positive) and number of infected (mCherry positive) were counted, and percent cell infection was computed.

### Cell viability assay

BMDMs were seeded at a density of 1×10^4^ cells/well in 96-well plates and infected with mCherry-*C. burnetii* at an MOI of 100 GE/cell as described above. At 6 and 12 DPI, cells were stained with Hoechst 33342 (2’-[4-ethoxyphenyl]-5-[4-methyl-1-piperazinyl]-2,5’-bi-1H-benzimidazole) (Biorad 1351304) and SYTOX green nucleic acid stain (Invitrogen/ThermoFisher S7020). Using a Molecular Devices ImageXpress Micro Confocal, total number of cells (Hoechst positive) and number of dead cells (SYTOX positive) were counted, and percent cell death was computed.

### Quantitative real-time reverse transcription PCR (qRT-PCR)

Total RNA was isolated from cells lysed in TRIzol (Invitrogen/ThermoFisher 15596026) using the Direct-Zol RNA miniprep kit (Zymo research R2052) and cDNA was synthesized using iScript Reverse Transcription Supermix (BioRad 1708840). qRT-qPCR was performed in triplicate using single tube TaqMan assay (Invitrogen/ThermoFisher; Mm00443258_m1, Mm99999915_g1). For all gene expression data, *Gapdh* was used as an endogenous normalization control.

### Enzyme-linked immunosorbent assay (ELISA)

BMDMs were seeded at a density of 2×10^5^ cells/well in 12-well plates and infected with mCherry-*C. burnetii* at an MOI of 300 GE/cell as described above. At 12 DPI, cell-free supernatants were collected from mock- and *C. burnetii*-infected cells. Supernatants were diluted with assay buffer and analyzed for presence of TNFα by ELISA (Invitrogen/ThermoFisher BMS607-3 or BMS607-2HS) according to manufacturer’s protocol.

### Statistical analyses

Densitometry was performed using ImageJ software. Statistical analyses were completed using GraphPad Prism 9. Statistical tests performed are specified in figure captions. Results shown are representative of at least three biological replicates from at least two independent experiments, as specified in the figure captions. Unless otherwise indicated, all error bars represent the standard deviation.

## Author Contributions

Conceptualization by C. A. O. and A. G. G. Methodology by C. A. O., C. L., H. S. K., and A. G. G. Reagents provided by H. S. K. and A. G. G. Experiments, optimization, and statistics completed by C. A. O., C. L., and N. H. Figures made by C. A. O. in consultation with A. G. G. Writing by C. A. O. and revised by A. G. G., C. L., N. H., and H. S. K.

## Acknowledgments

The *C. burnetii* clone 4 RSA439 used in this study was a gift from Dr. Robert A. Heinzen (Rocky Mountain Laboratories, NIH, Hamilton, MT). We thank Dr. Hongyan Guo and lab for supplying mutant mouse femurs for BMDM isolation, Dr. Santanu Bose and lab for supplying THP-1 cells, Miyoung Lee for experimental support, and Dr. Manish Chauhan for experimental support and technical expertise. This research was supported by NIH / National Institute of Allergy and Infectious Diseases (NIAID) grant R01 AI139051 to A. G. G and by a Poncin Fellowship to C.A.O.

## Ethics Declarations

The authors declare no competing interests.

